# Intra-articular AAV-PHP.S mediated chemogenetic targeting of knee-innervating dorsal root ganglion neurons alleviates inflammatory pain in mice

**DOI:** 10.1101/2020.02.08.939066

**Authors:** Sampurna Chakrabarti, Luke A. Pattison, Balint Doleschall, Rebecca H. Rickman, Helen Blake, Gerard Callejo, Paul A. Heppenstall, Ewan St. John Smith

## Abstract

**Objective:** Joint pain is the major clinical symptom of arthritis that affects millions of people. Controlling the excitability of knee-innervating dorsal root ganglion (DRG) neurons (knee neurons) could potentially provide pain relief. Therefore, our objective was to evaluate whether the newly engineered adeno-associated virus (AAV) serotype, AAV-PHP.S, can deliver functional artificial receptors to control knee neuron excitability following intra-articular knee injection.

**Methods:** AAV-PHP.S virus packaged with dTomato fluorescent protein and either excitatory (G_q_) or inhibitory (G_i_) designer receptors activated by designer drugs (DREADDs) was injected into the knee joint of adult mice. Labelling of DRG neurons by AAV-PHP.S from the knee was evaluated using immunohistochemistry. Functionality of G_q_- and G_i_-DREADDs was evaluated using whole-cell patch clamp electrophysiology on acutely cultured DRG neurons. Pain behavior in mice was assessed using a digging assay, dynamic weight bearing and rotarod, before and after intra-peritoneal administration of the DREADD activator, Compound 21.

**Results:** We show that AAV-PHP.S can deliver functional genes into the DRG neurons when injected into the knee joint in a similar manner to the well-established retrograde tracer, fast blue. Short-term activation of AAV-PHP.S delivered Gq-DREADD increases excitability of knee neurons in vitro, without inducing overt pain in mice when activated in vivo. By contrast, in vivo G_i_-DREADD activation alleviated complete Freund’s adjuvant mediated knee inflammation-induced deficits in digging behavior, with a concomitant decrease in knee neuron excitability observed in vitro.

**Conclusions:** We describe an AAV-mediated chemogenetic approach to specifically control joint pain, which may be utilized in translational arthritic pain research.

## Introduction

Peripheral sensitization, manifested by an increase in the excitability of dorsal root ganglion (DRG) neurons, underlies many chronic pain pathologies, such as inflammatory arthritis (1). DRG neurons display great heterogeneity, based upon both gene expression (2,3) and functional attributes (4), and this heterogeneity is further compounded by target innervation (5,6). The variation in DRG neurons offers a unique opportunity to selectively tune the excitability of a distinct subset of DRG neurons in order to provide pain relief with reduced side-effects. For example, we have recently shown that the excitability of knee-innervating DRG neurons (identified by retrograde tracing) is increased in a mouse model of inflammatory joint pain (7) and after incubation with human osteoarthritis (OA) synovial fluid samples (8). These results suggest that modulation of the knee-innervating DRG neuron subset (knee neurons) could help control arthritic pain.

One way of modulating neuronal excitability is to induce expression of inhibitory or excitatory designer receptors exclusively activated by exogenous chemical actuators which by themselves do not have any endogenous effects. Modified muscarinic receptor based designer receptors activated by designer drugs (DREADDs) is a technology based on this principle that can increase or decrease neuronal (mostly in the central nervous system, CNS) firing, which consequently affects a variety of behaviors (reviewed in (9,10)), such as enhanced feeding (11) or decreased wakefulness (12). In the peripheral nervous system (PNS), activation of the inhibitory DREADD hM_4_D(G_i_) in voltage-gated sodium channel 1.8 (Na_V_1.8) expressing DRG neurons decreased knee hyperalgesia and mechanical allodynia, along with a decrease in DRG neuron firing, in mice with early experimental OA pain induced by surgical destabilization of the medial meniscus (13). This attenuation of hyperalgesia was at a similar level when compared to administration of 10 mg/kg morphine, thus suggesting that peripherally acting analgesics can be comparably potent to well-established opioids. Similarly, activating G_i_-DREADD in transient receptor potential vanilloid 1 (TRPV1) expressing DRG neurons (14) increased the heat pain threshold and reduced neuronal excitability in mice. Both of these studies used transgenic mice and therefore are less translatable across species due to the technical difficulties associated with targeting sensory neuron sub-populations in species without such transgenic tractability. In wild-type mice, G_i_-DREADD delivered intraneurally to the sciatic nerve via adeno-associated virus 6 (AAV6) was able to increase both mechanical and thermal thresholds (15). Importantly however, none of these studies were specific to DRG neuron subsets innervating specific organs.

AAVs are useful tools for gene transfer that have been used for gene therapy in a variety of human diseases (16), with multiple AAV-based clinical trials currently underway for arthritis (Clinical trial # NCT02727764, NCT02790723 (17)). AAVs can be utilized in conjunction with DREADD technology to selectively modulate neuronal activity of specific neuronal circuitry. Indeed, this has been achieved in the CNS (18). However, delivering genes by AAV injection into a peripheral organ to DRG neurons is challenging because of the low transduction capability of AAVs and the large anatomical distances involved in the PNS (19). A variety of AAV serotypes have shown little efficacy in transducing DRG neurons when injected subcutaneously, intra-muscularly or intra-plantarly in adult mice (20,21). To date direct injection into DRG (22) or intrathecal injection (23,24) are the best ways for transducing DRG neurons, however, these routes of administration are invasive, technically complicated to perform and do not enable transduction of neurons innervating a defined target. In the present study, we provide evidence that the PNS specific AAV serotype, AAV-PHP.S (19), can infect DRG neurons with functional cargo following injection into the knee joint. Furthermore, using the inhibitory DREADD, hM_4_D(G_i_), as a cargo, we show that its activation normalizes the inflammatory pain induced deficit in digging behavior in mice, which is an ethologically relevant spontaneous pain measure indicating well-being (25). This study thus extends the use of AAV and DREADD technologies to study DRG neurons infected from a peripheral organ, which can have future clinically-relevant applications in controlling pain pathologies.

## Materials and Methods

### Animals

10-15 week old C57BL/6J (Envigo) mice (n = 30) of both sexes were used in this study. Mice were housed in groups of up to 5 in a temperature controlled (21 °C) room on a 12-hour/light dark cycle with food, water and appropriate enrichment available *ad libitum*. Mice used in this study were regulated under the Animals (Scientific Procedure) Act 1986, Amendment Regulations 2012. All protocols were approved by a UK Home Office project license (P7EBFC1B1) and reviewed by the University of Cambridge Animal Welfare and Ethical Review Body.

### Viruses

AAV-PHP.S-CAG-tdTomato (Addgene #59462-PHP.S) was purchased from Addgene. AAV plasmids for DREADD viruses were purchased from Addgene (Table S1) and packaged at the European Molecular Biology Laboratory, Rome as has been described previously (26). Briefly, ten 150 mm dishes of HEK293T (ATCC) cells were triple transfected with plasmids (Table S1) of AAV-PHP.S, helper (Agilent) and cargo in a 1:4:1 ratio with PEI reagent (1:3 plasmid to PEI ratio, Sigma). Three days after transfection, cells and media were collected and this mixture was centrifuged at 3700xg at 4°C to remove debris and then concentrated by ultrafiltration using Vivaflow 200 (Sartorius). The purified AAV particles were collected after running the viral concentrate through an iodixanol column (Opti-Prep density gradient medium, Alere Technologies) by ultracentrifugation (Beckmann, L8-70M) at 44400xg for 2 hr at 18 °C followed by filtration using a 100 kDa filter to further concentrate the sample and for buffer exchange. Viral titres (vg/ml) were measured by probing for WPRE regions (forward: GGCTGTTGGGCACTGACAAT, reverse: CCGAAGGGACGTAGCAGAAG) using SyBR green qPCR of linearized virus particles as described previously (26) using a StepOnePlus Real Time PCR system, following the manufacturer’s guidelines on settings (Applied Biosystems).

### Knee injections

All knee injections were conducted under anesthesia (100 mg/kg ketamine and 10 mg/kg xylazine, intra-peritoneally) through the patellar tendon. For experiments with retrograde tracing, 1.5 μl of fast blue (FB) or titer-matched AAV-PHP.S viruses were injected intra-articularly into knee joints (∼4×10^11^ vg of AAV-PHP.S-CAG-dTomato and ∼5×10^11^ vg of AAV-PHP.S packaged with G_i_ and G_q_-DREADD into each knee). The titer values were chosen based on a pilot experiment conducted with AAV-PHP.S-CAG-YFP, injection of 2 ul (∼3×10^11^ vg) and 10 ul (∼10^^12^ vg) labelled a similar number of neurons as assessed by acute culture. For experiments involving inflammation, 7.5 μl CFA (10 mg/ml, Chondrex) was injected into the left knee and Vernier’s calipers were used to measure knee width (as before (7)) pre- and 24-hour post-CFA injection.

### Behavioral testing

All behavioral experiments were carried out between 10:00 and 13:00 in the presence of one male and one female experimenter. Mice were assigned randomly to control and experimental groups and at least two cohorts of mice (8-12 mice in each group) were assessed in each group on separate days. Mice were trained on digging and rotarod one day before the test days. The following groups were tested in this study:

1. **Compound 21 (C21) controls:** Behavioral tests (described below) were performed on mice with no knee injections 20 min before and after C21 (2 mg/kg diluted in sterile saline from a stock of 100 mM in ethanol, i.p., Tocris) injection.
2. **Activation of G**_**q**_**-DREADD:** 3-4 weeks after intra-articular administration of virus, mice were behaviorally tested 20 min before and after vehicle (1:100 ethanol in sterile saline) or C21 injections.
3. **Activation of G**_**i**_**-DREADD:** 3-4 weeks after intra-articular administration of virus, baseline behavioral tests were conducted on mice (pre-CFA). CFA was injected into the knee joints the next day to induce inflammation. 24-hours after that, animals were re-tested 20 min before (post-CFA) and after (post-C21) vehicle or C21 injections.

### Digging

Digging behavior was measured as an assessment of spontaneous pain as described before (7) for three min in a standard cage with a wire lid, filled with Aspen midi 8/20 wood chip bedding (LBS Biotechnology). For training, mice were habituated in the test room in their home cages for 30 min then they were allowed to dig twice for 3 min with 30 min break in between. On each subsequent test day, mice were habituated and tested once on the 3 min paradigm. Test sessions were video recorded from which the digging duration was later independently coded by the experimenters, blinded to the conditions. Number of dig sites (burrows) was coded on test days by the experimenters.

### Rotarod

Locomotor function and coordination of mice were tested using a rotarod apparatus (Ugo Basile 7650). Mice were tested on a constant speed rotarod at 7 rpm for 1 min, then in an accelerating program (7-40 rpm in 5 min) for 6 min. The same protocol was used to train mice one day before testing. Mice were removed from the rotarod after two passive rotations or when they fell from the rotarod. Mice were video recorded on test days and one experimenter blinded to the conditions coded these videos for latency(s) to passive rotation or fall.

### Dynamic weight bearing

Deficit in weight bearing capacity is a characteristic measure of spontaneous inflammatory pain behavior and we measured this behavior using a dynamic weight bearing (DWB) device (Bioseb) in freely moving mice for 3 min. Animals were not trained in this device and coding was done by one experimenter, blinded to the conditions. In at least 1 min 30 sec of the 3 min recording, fore- and hind paw prints were identified using the two highest confidence levels (based on correlation between manual software algorithm tracking) of the in-built software, at least 30 s of which was manually verified.

### DRG neuron culture

Lumbar DRG (L2-L5) were collected from a subset of mice in the vehicle group 3-4 weeks after virus injections in ice cold dissociation media (as before (7)). DRG were then enzymatically digested followed by mechanical trituration (7). Dissociated DRG neurons, thus isolated, were then plated onto poly-D-lysine and laminin coated glass bottomed dishes (MatTek, P35GC-1.5-14-C) in DRG culture medium contained L-15 Medium (1X) + GlutaMAX-l, 10% (v/v) fetal bovine serum, 24 mM NaHCO_3_, 38 mM glucose, 2% penicillin/streptomycin and maintained overnight (8-10-hours) in a humidified, 37 °C, 5% CO_2_ incubator before recording.

### Electrophysiology

For control experiments, DRG neurons were bathed in extracellular solution containing (ECS, in mM): NaCl (140), KCl (4), MgCl_2_ (1), CaCl_2_ (2), glucose (4) and HEPES (10) adjusted to pH 7.4 with NaOH while recording. To investigate effects of C21 activation, neurons were incubated 10 min prior to the start of recording and throughout the recordings in 10 nM C21 (diluted in ECS from 100 mM stock). Only virus transduced mice identified by their fluorescence upon excitation with a 572 nm LED (Cairn Research) were recorded using patch pipettes of 5–10 MΩ (P-97 Flaming/Brown puller, Sutter Instruments) containing intracellular solution (in mM): KCl (110), NaCl (10), MgCl_2_ (1), EGTA (1), HEPES (10), Na_2_ATP (2), Na_2_GTP (0.5) adjusted to pH 7.3 with KOH.

Action potentials (AP) were recorded in current clamp mode without current injection (to investigate spontaneous AP firing) or after step-wise injection of 80 ms current pulses from 0 – 1050 pA in 50 pA steps using a HEKA EPC-10 amplifier (Lambrecht) and the corresponding Patchmaster software. AP properties were analysed using Fitmaster software (HEKA) or IgorPro software (Wavemetrics) as described before (7). Neurons were excluded from analysis if they did not fire an AP in response to current injections.

### Immunohistochemistry

Mice with unilateral, co-injections of AAV-PHP.S-tdTomato (2 ul) and FB (1.5 ul) were transcardially perfused with 4% (w/v) paraformaldehyde (PFA; in PBS, pH 7.4) under terminal anesthesia (sodium pentobarbital, 200 mg/kg, i.p.). L2-L5 DRG were then collected from the injected side, while L3-L4 were collected from the non-injected side and post-fixed in Zamboni’s fixative for 1-hour, followed by overnight incubation in 30% (w/v) sucrose (in PBS) at 4 °C for cryo-protection. DRG were then embedded, snap-frozen, sectioned and stained as described previously (7) using anti-TRPV1 antibody (1:500, Alomone, AGP-118, anti-guinea pig polyclonal) with Alexa-488 conjugated secondary antibody (1:500, Jackson Laboratory, 706-545-148, anti-guinea pig). Positive neurons were scored as has been reported previously (7) using a R toolkit (https://github.com/amapruns/Immunohistochemistry_Analysis) followed by manual validation. Briefly, mean grey value (intensity) of all neurons on 1-3 sections from each DRG level of interest from each mouse was measured using ImageJ. A neuron was scored positive for a stain if it had an intensity value greater than average normalized minimum grey value across all sections + 2 times the standard deviation.

### Statistics

Comparisons between two groups were made using appropriate (paired for repeated measures and unpaired otherwise) Student’s t-test while three group comparisons were made using one-way analysis of variance (ANOVA) followed by Holm-Sidak multiple comparison tests. Chi-sq tests were utilized to compare proportions of categorical variables. Data are presented as mean ± standard error of mean.

## Results

### 1. Knee injected AAV-PHP.S robustly transduces DRG neurons

AAV-PHP.S was engineered to have a higher specificity towards peripheral neurons (19) and thus we hypothesized that this AAV serotype would be able to transduce DRG neurons when injected intra-articularly into the knee joint. Consistent with our hypothesis, when AAV-PHP.S-CAG-dTomato and the commonly used retrograde tracer fast blue (FB) were co-injected unilaterally into one knee of mice (n = 3, female), we observed both FB and virus labelling (Fig 1A). In agreement with previous reports using FB and other retrograde tracers (7,27,28), we observed a similar proportion of labelling in the lumbar DRG with FB and AAV-PHP.S-CAG-dTomato (Fig 1B). Across L2-L5 DRG, there was ∼ 40 % co-labelling of neurons with FB and AAV-PHP.S-CAG-dTomato suggesting that neither retrograde tracer is able to label the entire knee neuron population (Fig 1C, Fig S1). However, area analysis of the labelled neurons suggests that similar sized neurons are targeted by both strategies (Fig S1). Furthermore, we observed minimum labelling in the contralateral side (Fig S1). Previous reports suggest that ∼40 % of knee neurons are TRPV1 expressing putative nociceptors (7,28). Similarly, with immunohistochemistry analysis of DRG neurons, we find that ∼30 % of viral labelled and FB labelled neurons (mostly small diameter neurons) express TRPV1 (Fig 1D, Fig S1) suggesting that viral transduction did not substantially alter expression of the nociceptive protein, TRPV1. Taken together, we find that intra-articular injection of AAV-PHP.S-CAG-dTomato in the knee joint transduces mouse DRG neurons in a similar manner to a routinely used retrograde tracer.

**Fig. 1.**
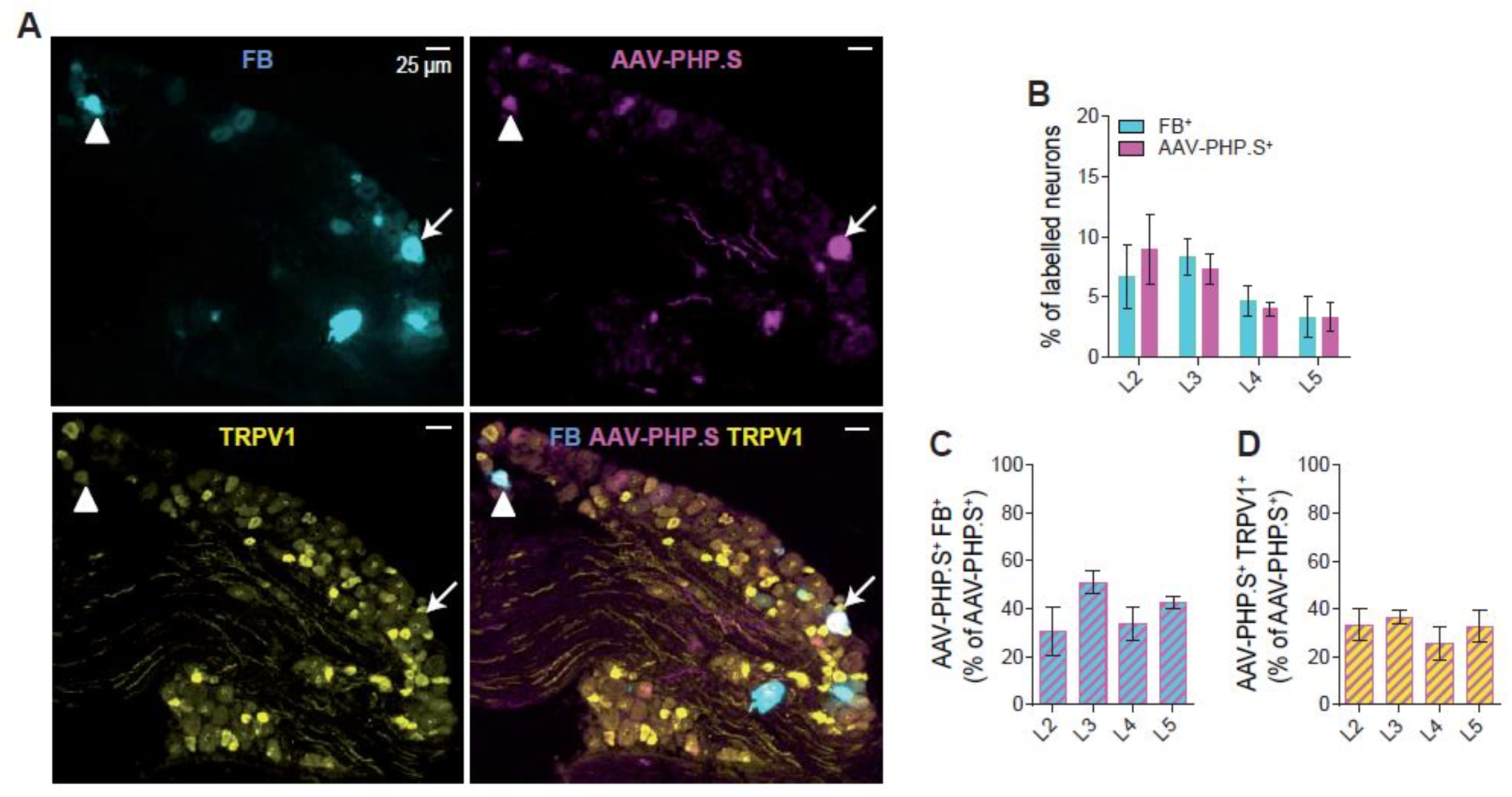
Retrograde tracing of knee-innervating DRG neurons using FB and AAV-PHP.S-CAG-dTomato. A) Representative images of a whole L3 DRG section showing knee neurons traced using FB (blue), AAV-PHP.S-CAG-dTomato (pink) and stained using an anti-TRPV1 antibody (yellow). White triangle is pointing at a neuron that had FB, AAV-PHP.S and TRPV1 colocalization, while the white arrow is pointing at a neuron that shows only FB and AAV-PHP.S co-localization. B) Bar graphs showing percent of labelled neurons in L2-L5 DRG with FB and AAV-PHP.S. Percent of neurons showing co-localization of FB and AAV-PHP.S (C), and TRPV1 and AAV-PHP.S (D), expressed as a percent of AAV-PHP.S^+^ neurons. The error bars represent the SEM in the data obtained from three female mice.

### 2. Excitatory G_q_-DREADD delivered intra-articularly by AAV-PHP.S-hSyn-hM_3_D(G_q_)-mCherry does not change spontaneous pain behavior, but provokes a deficit in motor coordination

Next we tested whether AAV-PHP.S can deliver functional hM_3_D(G_q_) cargo into knee neurons via intra-articular injection, using whole-cell patch clamp on acutely dissociated neurons isolated from mice with no previous exposure to the DREADD activator Compound (C21). C21 was chosen as the DREADD activator since it has good bioavailability (plasma concentration measured to be at 57% 1 hour post injection), is not converted into clozapine or clozapine-N-oxide (29) and had no off-target effect in the behavioral tests conducted in this study in naïve mice not injected with DREADDs (Fig S2). G_q_-DREADDs couple to G_q_PCR pathways and thus their activation causes neuronal excitation (9). Therefore, we hypothesized that when incubated with 10 nM C21, virally transduced neurons would be hyperexcitable compared to virally transduced neurons bathed in normal extracellular solution (ECS). In agreement with this hypothesis, we observed an increased number of mostly small and medium diameter (Fig S3) mostly nociceptive (characterized by having an AP half peak duration > 1 ms and with a “hump” during repolarization, Fig S3, (30)) neurons (Ctrl, 5.3% vs. C21, 27.8% p = 0.03, chi-sq test) firing action potentials (AP) without injection of current in the C21 group (Fig 2A,B, S3). Moreover, upon injection of increasing stepwise current injections, the action potential (AP) threshold was decreased (p = 0.02, unpaired t-test) in the C21 group (Fig 2C); no change was observed in other electrical properties measured in these neurons (Table 1). Our data also suggest that virally transduced neurons are viable because the reported AP threshold is very similar to what we have observed previously in FB+ knee-innervating neurons (31).

**Table 1:** Action potential properties of G_q_-DREADD intra-articularly transduced knee neurons. RMP = Resting membrane potential, HPD = Half peak duration. * = p < 0.05 unpaired t-test.

|  | **Ctrl** |  | **C21** |  |
| --- | --- | --- | --- | --- |
|  | **(n = 19)** |  | **(n = 18)** |  |
|  | Mean | SEM | Mean | SEM |
| **RMP (mV)** | -40.8 | 3.0 | -42.7 | 2.8 |
| **Threshold (pA)** | 415.8 | 69.7 | 211.1* | 51.4 |
| **HPD (ms)** | 2.6 | 0.5 | 1.9 | 0.3 |
| **Afterhyperpolarization**<br>**amplitude (AHP peak,**<br>**mV)** | 14.7 | 2.1 | 16.8 | 2.8 |

**Fig. 2.**
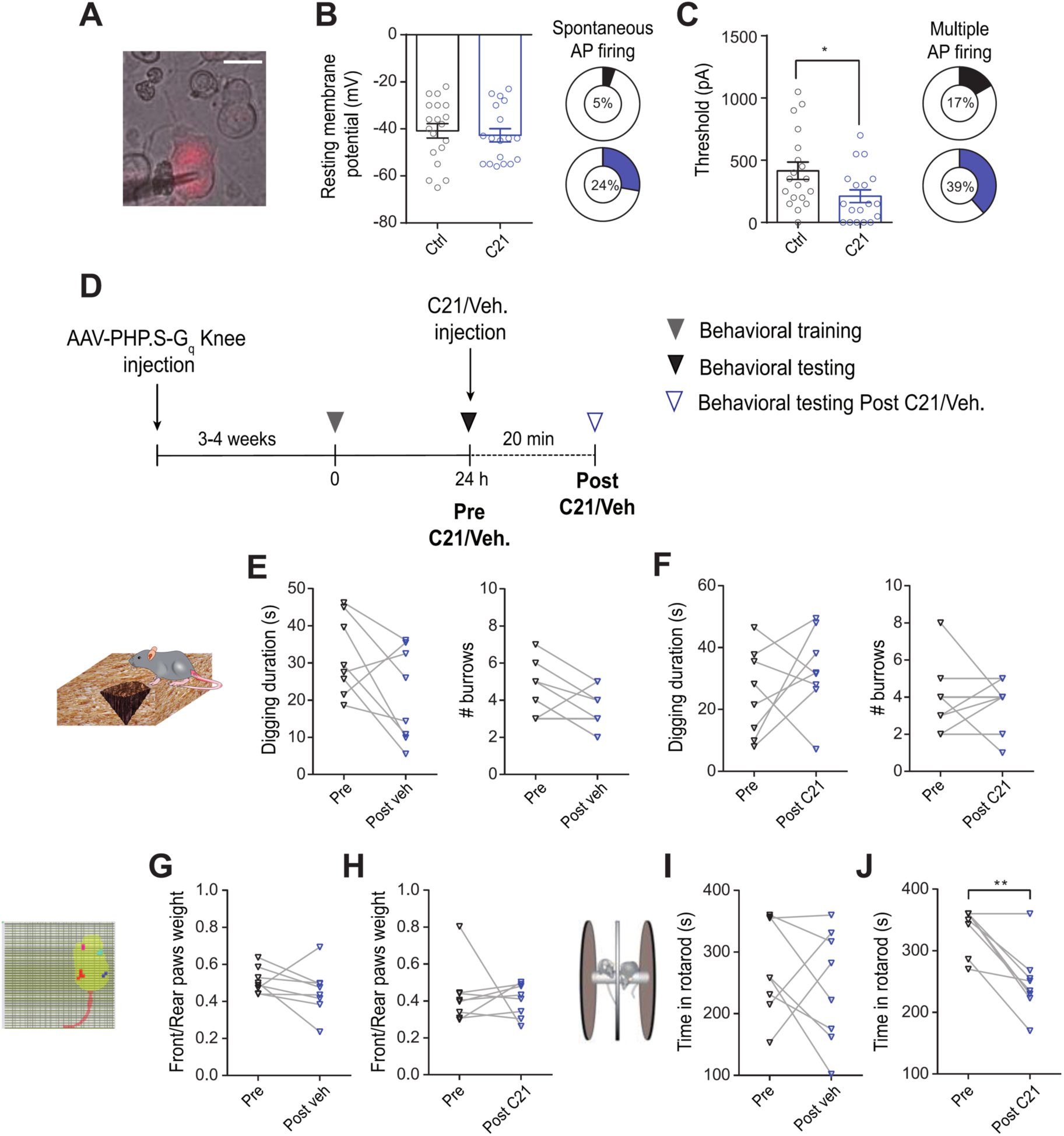
G_q_-DREADD activation of knee neurons in-vitro and in-vivo. A) Representative image of an AAV-PHP.S-hSyn-hM_3_D(G_q_)-mCherry transduced neuron (Scale bar = 25 µm, triangular shadow = recording electrode). B) bars showing resting membrane potential in Ctrl (black, n = 19) and C21 (blue, n = 18) conditions, pie-chart showing percent of neurons in each condition that fired spontaneous AP. C) Bars showing AP firing threshold in Ctrl (black, n = 19) and C21 (blue, n = 18) conditions, pie-chart showing percent of neurons in each condition that fired multiple AP upon current injection. *p < 0.05, unpaired t-test. Data obtained from 3 females and 2 male mice. D) Timeline showing when behaviors were conducted. Digging duration and burrows (along with schematic diagram), ratio of front and rear paw weight (along with schematic diagram) and rotarod behavior (along with schematic diagram) before and after vehicle (E, G, I) or C21 injection (F, H, J). ** p < 0.01, paired t-test. Data obtained from 4 female and 4 male mice. Error bars = SEM.

Based on previous studies (7,8), we hypothesized that the measured increased excitability of knee-innervating neurons in vitro via the G_q_-DREADD system would cause pain-like behavior in mice in vivo, which was tested by measuring digging behavior (a measure of spontaneous pain as described previously (7,30)), dynamic weight bearing and rotarod behavior (a measure of motor co-ordination) (31) (Timeline in Fig 2D). Three weeks after virus injection into both knee joints, mice (n = 8, 4 males, 4 females in each group) injected with vehicle or C21 did not show changes in digging behavior (Digging duration: Pre vs. Post veh, 31.6 ± 3.7 ms vs. 21.3 ± 4.4 ms, Pre vs. Post C21, 25.2 ± 4.9 ms vs. 32.7 ± 4.7 ms; Number of burrows: Pre vs. Post veh, 4.6 ± 0.5 vs. 3.5 ± 0.4, Pre vs. Post C21, 3.9 ± 0.7 vs. 3.6 ± 0.5, Fig 2E,F) or weight bearing (Front/Rear paw weight ratio: Pre vs. Post veh, 0.5 ± 0.02 vs. 0.5 ± 0.04, Pre vs. Post C21, 0.4± 0.06 vs. 0.4 ± 0.03, Fig 2G,H). By contrast, after injection of C21, mice showed a marked decline in their ability to remain on the rotarod (Pre vs. Post veh, 273.5 ± 27.2 s vs. 243.9 ± 32.7 s, Pre vs. Post C21, 336.1 ± 12.9 s vs. 249.0 ± 18.9 s, p = 0.002, paired t-test) suggesting a deficit in their motor co-ordination (Fig 2I,J). DRG for all virus injected mice were visualized under a fluorescence microscope to check for viral transduction and a subset of these DRG were further analyzed to reveal similar transduction profiles between AAV-PHP.S-hSyn-G_q_DREADD-mCherry and AAV-PHP.S-CAG-dTomato (Fig S3), i.e. different promoters do not significantly affect transduction.

Taken together, our data show that AAV-PHP.S delivers functional G_q_-DREADD into knee neurons and that when virally transduced knee neurons are activated they do not produce overt pain-like behavior in vivo, but do cause a deficit in motor coordination.

### 3. Inhibitory G_i_-DREADD delivered intra-articularly by AAV-PHP.S-hSyn-hM_4_D(G_i_)-mCherry reverses digging behavior deficits associated with inflammatory pain

Intra-articular injection of complete Freund’s adjuvant (CFA) induces robust knee inflammation in mice (Ctrl knee: pre CFA day, 3.1 ± 0.03 mm, post CFA day, 3.1 ± 0.03 mm; CFA knee: pre CFA day, 3.1 ± 0.02 mm, post CFA day, 4.0 ± 0.05 mm, n = 24, p < 0.0001, paired t-test, Fig 3A,B) and has been previously shown to increase the excitability of knee neurons innervating the inflamed knee compared to those innervating the contralateral side (7). The post-CFA knee measurements were conducted at the end of behavioral measurements, thus suggesting that regardless of G_i_-DREADD activation knee inflammation persisted at 24-hour post-CFA injection.

**Fig. 3.**
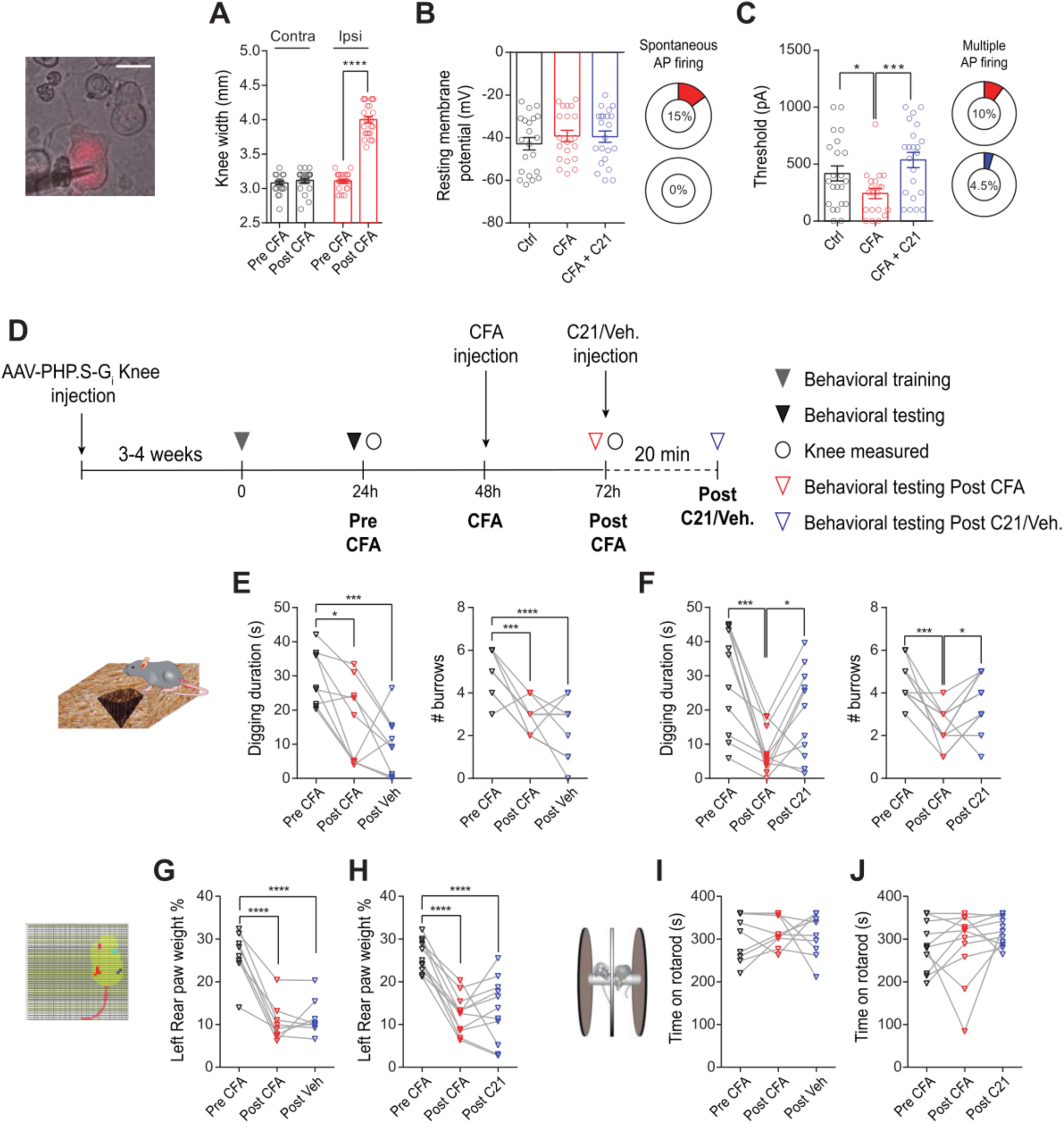
G_i_-DREADD activation of knee neurons in vitro and in vivo. A) Bars representing knee width before and after CFA injection in the non-injected (contra, black) and injected knee (ipsi, red), **** p < 0.0001, n = 20, paired t-test. B) Bars showing resting membrane potential in Ctrl (black, n = 22), CFA (red, n = 20) and CFA+C21 (blue, n = 22) conditions, pie-chart showing percent of neurons in CFA and CFA+C21 condition that fired spontaneous AP. C) Bars showing AP firing threshold in Ctrl (black, n = 22), CFA (red, n = 20) and CFA+C21 (blue, n = 22) conditions, pie-chart showing percent of neurons in CFA and CFA+C21 condition that fired multiple AP upon current injection. *p < 0.05, ***p < 0.001 ANOVA and Holm-Sidak multiple comparison test. Data obtained from 2 female and 2 male mice. D) Timeline showing when behaviors were conducted. Digging duration and number of burrows, rear left paw weight expressed as percent of body weight and rotarod behavior pre and post-CFA and after vehicle (E, G, I, 4 female and 5 male mice) or C21 injection (F, H, J, 4 female and 7 male mice). *p < 0.05, ** p < 0.01, ***p < 0.001, **** p < 0.0001 repeated measures ANOVA and Holm-Sidak multiple comparison test. Error bars = SEM.

We hypothesized that incubating G_i_-DREADD expressing knee neurons from the CFA side with C21 would reverse this increased neuronal excitability. Using whole-cell patch clamp electrophysiology of small and medium diameter (Fig S4) knee neurons, although there was no change in the resting membrane potential across any condition (Fig 3C, Table 2), the percentage of CFA knee neurons firing spontaneous AP decreased after G_i_-DREADD activation (CFA, 15% vs. CFA+ C21, 0%, p = 0.02, chi-sq test, Fig 3C,S3). Moreover, in the absence of G_i_-DREADD activation, CFA knee neurons had a decreased AP threshold compared to neurons from the control side, but the AP threshold of CFA knee neurons that were incubated in C21 was similar to that of neurons from the control side (p = 0.005, ANOVA, Fig 3D). These results suggest that G_i_-DREADD activation reverses CFA-induced increase in nociceptor (based upon the criteria mentioned above, Fig S4) excitability in vitro. Other electrical properties between groups were unchanged (Table 2).

**Table 2:**
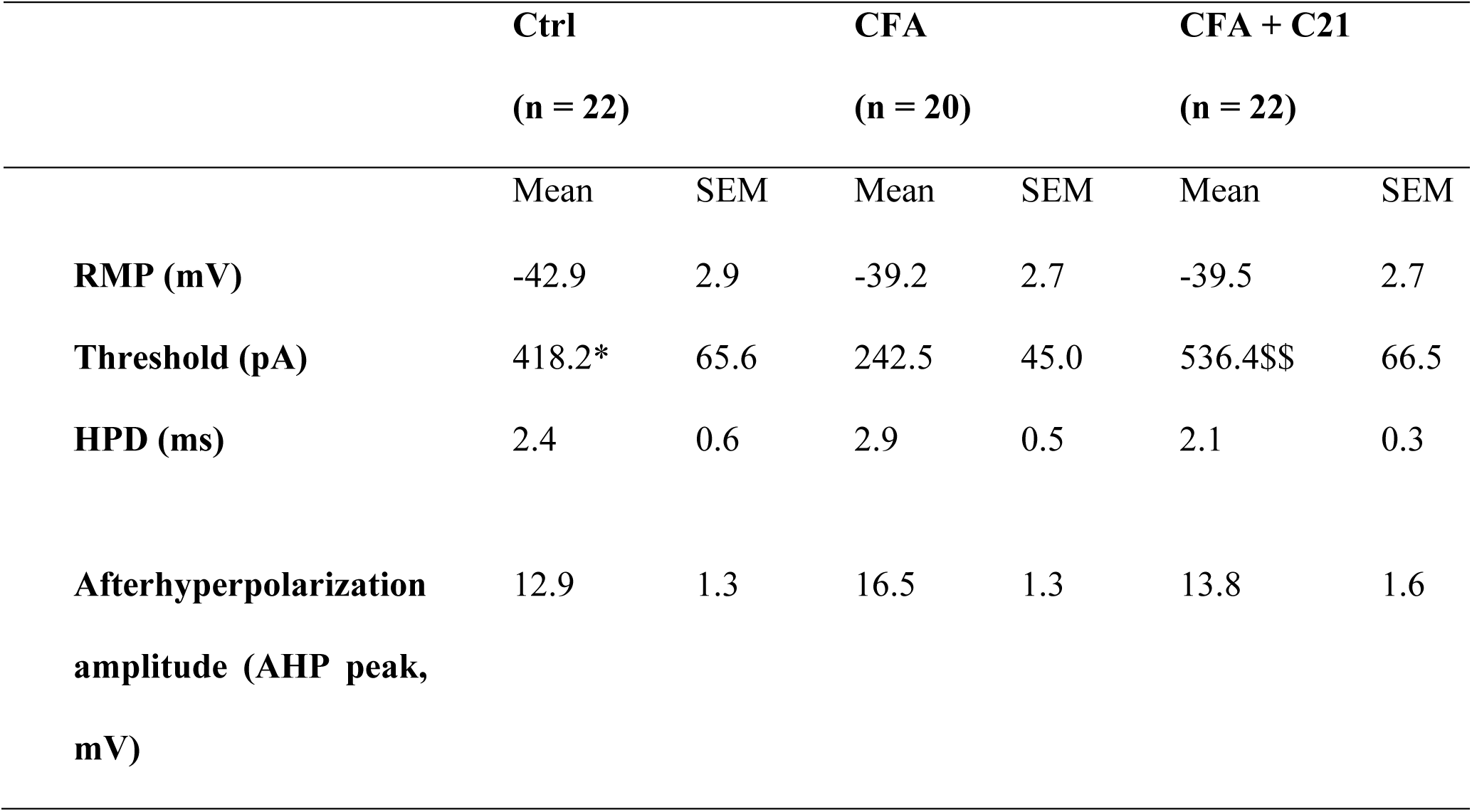
Action potential properties of G_i_-DREADD intra-articularly transduced knee neurons. RMP = resting membrane potential, HPD = Half peak duration. * = p < 0.05 CFA vs. Ctrl, $$ = p < 0.01 CFA vs. CFA+C21, unpaired t-test.

|  | **Ctrl** |  | **CFA** |  | **CFA + C21** |  |
| --- | --- | --- | --- | --- | --- | --- |
|  | **(n = 22)** |  | **(n = 20)** |  | **(n = 22)** |  |
|  | Mean | SEM | Mean | SEM | Mean | SEM |
| **RMP (mV)** | -42.9 | 2.9 | -39.2 | 2.7 | -39.5 | 2.7 |
| **Threshold (pA)** | 418.2* | 65.6 | 242.5 | 45.0 | 536.4\$\$ | 66.5 |
| **HPD (ms)** | 2.4 | 0.6 | 2.9 | 0.5 | 2.1 | 0.3 |
| **Afterhyperpolarization**<br>**amplitude (AHP peak,**<br>**mV)** | 12.9 | 1.3 | 16.5 | 1.3 | 13.8 | 1.6 |

The ability of G_i_-DREADD to modulate pain behavior in DRG neurons is unclear with one study showing an increase in latency to both thermal and mechanical stimuli (15), but another showing only an increase in the paw withdrawal latency to thermal stimuli (14). Nevertheless, based on the in vitro results in these studies, we hypothesized that activation of G_i_-DREADDs in knee neurons post CFA would reverse spontaneous pain behavior in mice (timeline in Fig 3E). In the control cohort of mice (n = 9, 5 males, 4 females) that received vehicle 24-hours after CFA injection, the CFA-induced decrease in digging behavior persisted compared to pre CFA (Digging duration: Pre CFA, 29.6 ± 2.7 ms, post CFA 16.6 ± 4.0 ms, post veh, 9.8 ± 2.8 ms, p = 0.0005, repeated measures ANOVA; Number of burrows: Pre CFA, 4.8 ± 0.3, post CFA 2.9 ± 0.3 ms, post veh, 2.3 ± 0.5 ms, n = 9, p < 0.0001, repeated measures ANOVA, Fig 3F). However, when C21 was administered to a separate cohort of mice (n = 11, 7 males, 4 females) 24-hours after CFA injection, there was a marked recovery in digging behavior (Digging duration: Pre CFA, 29.7 ± 4.5 ms, post CFA 7.8 ± 1.9 ms, post C21, 19.0 ± 3.9 ms, p = 0.0002, repeated measures ANOVA; Number of burrows: Pre CFA, 4.5 ± 0.3, post CFA 2.4 ± 0.3 ms, post C21, 3.6 ± 1.4 ms, p = 0.0005, repeated measures ANOVA, Fig 3G) suggesting that decreasing the excitability of knee neurons via G_i_-DREADD reduces inflammation induced spontaneous pain that is associated with an increase in the feeling of well-being demonstrated by more digging. In contrast, acute chemogenetic inhibition of knee neurons was insufficient to reverse the CFA-induced deficit in dynamic weight bearing (Rear left weight bearing as % of body weight: Pre CFA, 26.2 ± 2.0, post CFA, 10.6 ± 1.7, post veh, 11.6 ± 1.5, p < 0.0001; Pre CFA, 26.1 ± 1.1, post CFA, 12.4 ± 1.4, post C21, 13.3 ± 2.2, p < 0.0001, repeated measures ANOVA, Fig 3H,I) which might be because gait changes relating to weight bearing is more reflective of changes in joint biomechanics that are difficult to reverse by analgesics (32). Furthermore, no change in rotarod behavior was observed following CFA-induced knee inflammation suggesting that this model does not cause an overt change in gross motor function and similarly G_i_-DREADD activation also had no effect (Pre CFA, 303.8 ± 18.3 s, post CFA, 315.9 ± 11.9 s, post veh, 306.6 ± 16.8 s; Pre CFA, 282.7 ± 17.3 s, post CFA, 286.4 ± 25.3 s, post C21, 317.4 ± 10.3 s, Fig 3J,K). DRG for all virus injected mice were visualized under a fluorescence microscope to check for viral transduction (Fig S4).

Taken together, our results suggest that specific inhibition of knee neuron excitability can reverse inflammation-induced deficit in digging behavior.

## Discussion

The findings in this study show that the AAV-PHP.S serotype can retrogradely deliver cargo to DRG neurons in a peripheral tissue-specific manner when injected into the knee joints without the need for invasive procedures or the requirement to generate transgenic mice. The transduction efficacy of the virus is similar to the widely used retrograde tracer, FB. In-line with other co-labelling studies (33), the ∼40% co-localization of AAV-PHP.S and FB fluorescence suggests that not all knee neurons are labeled by either tracer and the less than 100% co-labeling is possibly due to their differing modes of retrograde transfer (34,35). Furthermore, we report that ∼30% of labelled neurons are TRPV1+, which fits with the previously reported identity of knee-innervating neurons as a subset being ∼39% TRPV1+, ∼53% CGRP+ and largely IB4 non-binding (28,39).

Using this system, we show that it is possible to increase or decrease knee neuron excitability in vitro when G_q_ or G_i_-DREADD cargoes were delivered by AAV-PHP.S respectively and hence provide joint specific pain control. This result can also be extended to study the role of anatomically specific neuronal excitability when exposed to a variety of stimuli or pharmacological interventions. In vivo, we restricted our chemogenetic activation to a short duration to reflect acute pain and within this timeframe saw no spontaneous pain-like behavior with activation of G_q_-DREADD in knee neurons. We surmise that G_q_-mediated sub-threshold activation (36) of the relatively low percentage of DRG neurons did not provide sufficient nociceptive input to drive change in ethologically relevant pain behavior; whereas the observed decrease in coordination suggests that we have behaviorally engaged the virally transduced neurons. Furthermore, we note that intra-articular injection described here would transduce DREADDs to both nociceptive and non-nociceptive population of knee neurons, therefore, a clear nocifensive behavior might not be apparent, i.e. a limitation of this study is that distinct subpopulations of knee neurons are not targeted, but future studies could address this once such populations have been described for the knee as has already been conducted for the colon (5). Future studies using a repeated dosing strategy could also be employed in our system for modelling chronic pain, being cautious of the potential risk of receptor desensitization (9).

Perhaps more relevant to future clinical applications in arthritic pain is the ability of G_i_-DREADDs to reverse pain behavior by decreasing neuronal excitability of knee neurons. Indeed, we show that G_i_-DREADD activation restores a deficit in digging behavior induced by inflammatory knee pain, similar to previous reports demonstrating normalization of burrowing/digging by non-steroidal anti-inflammatory drugs (41,42) and a peripherally restricted TRPV1 antagonist (7) after joint pain induced depression of this behavior. This strategy can be further refined to selectively inhibit genetically specific subpopulation of knee neurons by combining Cre-inducible viruses with their corresponding Cre-expressing transgenic mouse lines, and hence provide insights into relative contributions of different knee neuron sub-populations in arthritic pain. Selectively exciting specific knee neuron sub-populations with G_q_-DREADD might also produce pain-like behaviors that were not observed in this study. We also report that the CFA-induced deficit in weight bearing was not reversed following activation of G_i_-DREADD in knee neurons, consistent with a previous report observing that reversal of deficits in gait changes are difficult to achieve with analgesics (32).

Although findings from this study imply that modulating excitability of anatomically specific peripheral neurons could control arthritic pain, a number of challenges remain to be addressed before their clinical translation. Since virus transduction and expression profile is different between non-human primates and rodents, the expression profile of AAV-PHP.S needs to be first validated in non-human primates (37). Additional work is also required to engineer more PNS specific AAVs and to optimize DREADDs (38) and their corresponding ligands (39) for increasing transduction efficiency and regulating dosing.

Overall, the present study provides initial proof-of-concept that peripheral tissue innervating DRG neurons can be specifically modulated by AAVs, opening the door to future studies on gene therapy in controlling arthritic pain.

## Acknowledgements

**General** Authors would like to thank: Morris Poeuw and Fernanda Castro de Reis for technical help with viral production. Julie Gautrey and Anne Desevre for technical help with mouse behavior. Dewi Safitri for graphic design.

## Funding

This study was supported by Versus Arthritis Project grants (RG 20930 and RG 21973) to E. St. J. S. and G.C. S.C. was supported by Gates Cambridge Trust scholarship, Cambridge Philosophical Society research fund, Department of Pharmacology (University of Cambridge) travel award and Corpus Christi College (University of Cambridge) travel award. L.A.P. was supported by the University of Cambridge BBSRC Doctoral Training Programme (BB/M011194/1). P.A.H and B.D. were supported by EMBL.

## Author contributions

S.C. designed experiments, collected and analyzed data and wrote the manuscript. L.A.P. collected and analyzed behavioral data and revised the manuscript. B.D. helped with virus production. R.H.R. and H.B. helped with immunohistochemistry analysis. G.C. helped with behavioral experiments and revised the manuscript. P.A.H. and E. St. J. S. designed experiments and revised the manuscript. All authors approved the final version of the manuscript.

## Competing interests

The authors declare no conflict of interest.

## Data and materials availability

Data presented in this article will be available in the Cambridge University Apollo repository (https://doi.org/10.17863/CAM.46171).

## Supplementary Materials

**Fig. S1.**
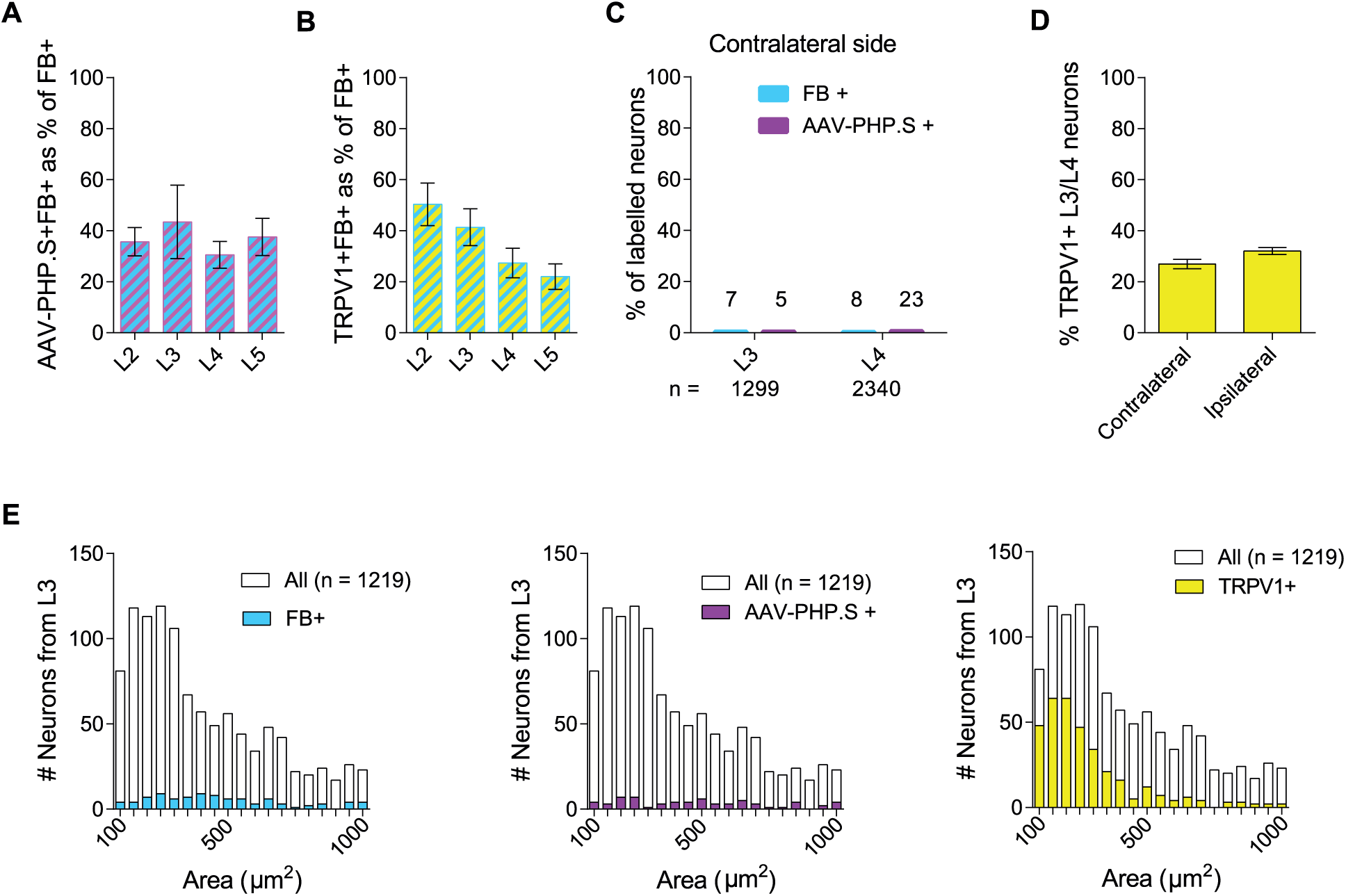
Specific Retrograde tracing of knee neurons with FB and AAV-PHP.S-CAG-dTomato. Bar graphs showing co-localization of FB and AAV-PHP.S-CAG-dTomato labelled neurons (A) and TRPV1 and AAV-PHP.S-CAG-dTomato labelled neurons (B) expressed as a percent of FB^+^ neurons. C) Percent of neurons showing FB and AAV-like fluorescence in the non-injected (contralateral) side. Numbers above bars represent total number of stained neurons and below the bars represent total number of neurons analyzed. D) Percent of TRPV1^+^ neurons in the injected (ipsilateral) and contralateral side. E) Histograms showing size distribution of L3 DRG neurons stained with FB, AAV-PHP.S and TRPV1 respectively. Data obtained from 3 female mice.

**Fig. S2.**
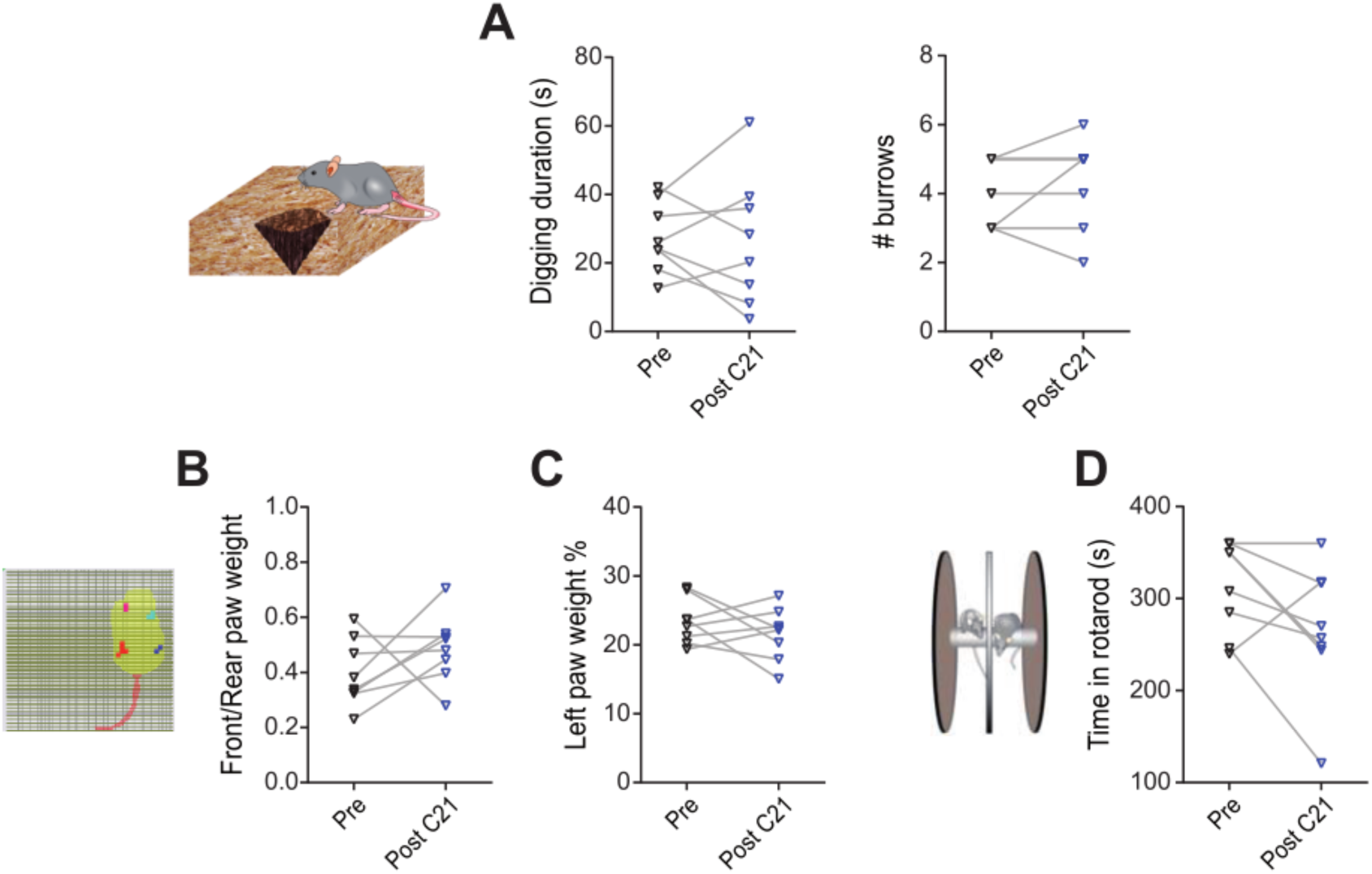
No off target behavioral effects of C21. Digging duration and number of burrows (A), ratio of front and rear paw weight (B), rear left paw weight expressed as a percent of body weight (C), and time on rotarod (D) measured before (black) and after (blue) C21 administration in mice with no virus injections. Data obtained from 4 female and 4 male mice.

**Fig. S3.**
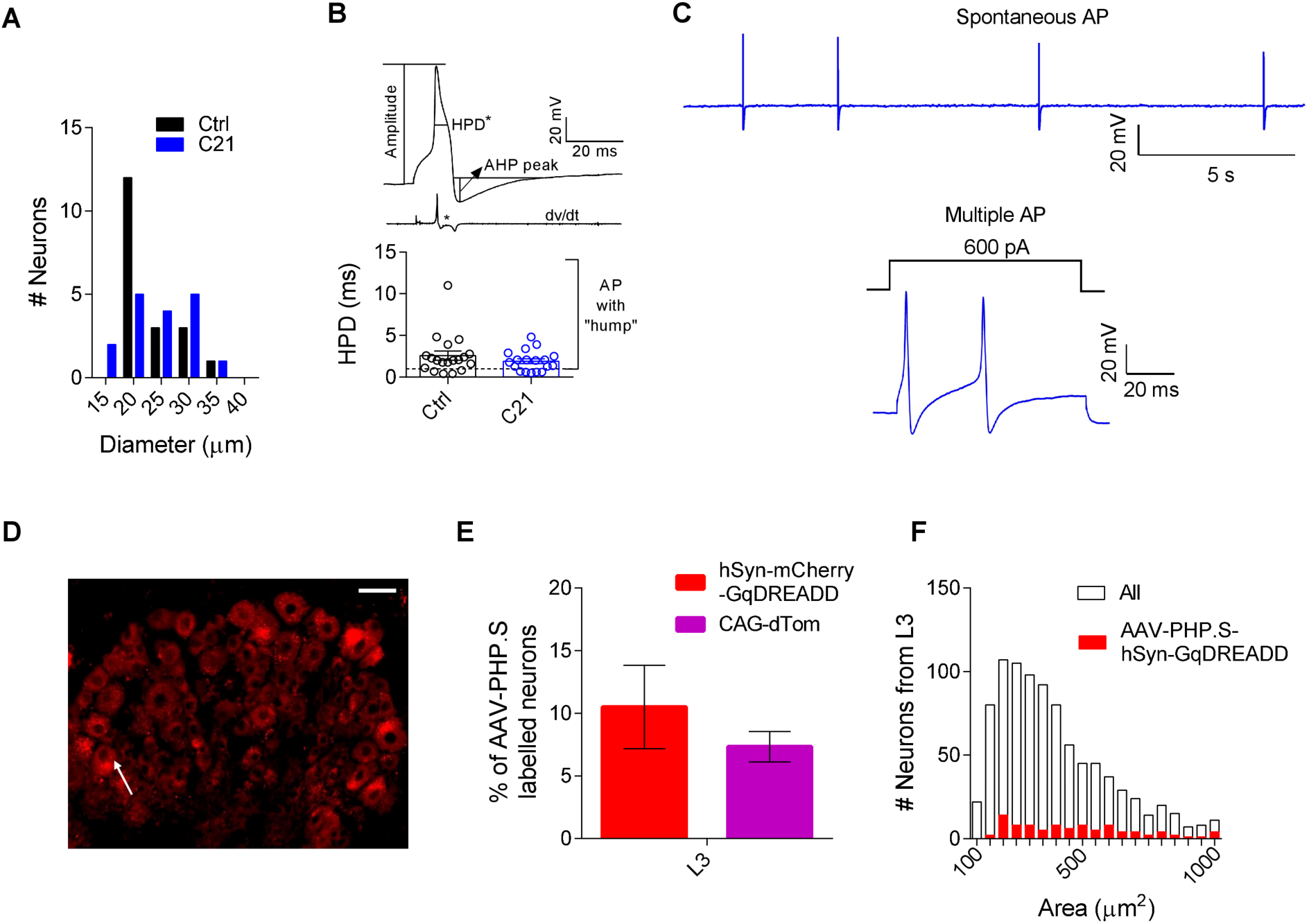
Profile of neurons transduced using AAV-PHP.S-hSyn-G_q_DREADD. A) Distribution of neuronal diameters transduced by GqDREADD and characterized using whole cell patch clamp electrophysiology. Data from 3 female and 2 male mice. B) Schematic diagram of an action potentials (AP) along with its dV/dt to highlight “hump” of a nociceptor (upper). Distribution of half peak duration (HPD) of AP recorded from transduced neurons. The black dotted horizontal line corresponds to HPD = 1 ms, thus demonstrating that most neurons had wide AP (lower). C) Representative traces of spontaneous and multiple firing of AP after C21 activation. D) Representative DRG (L3) section showing AAV-PHP.S-hSyn-G_q_DREADD labelled neuron (white arrow), scale bar = 50 μm. E) Percentage of labelled L3 neurons observed after AAV-PHP.S-hSyn-G_q_DREADD injection (data from 2 female and 2 male mice) compared to AAV-PHP.S-CAG-dTomato (data replotted from Fig 1). F) Histograms showing size distribution of L3 DRG neurons stained with AAV-PHP.S-hSyn-G_q_DREADD (n = 951, 3 female and 2 male mice).

**Fig S4.**
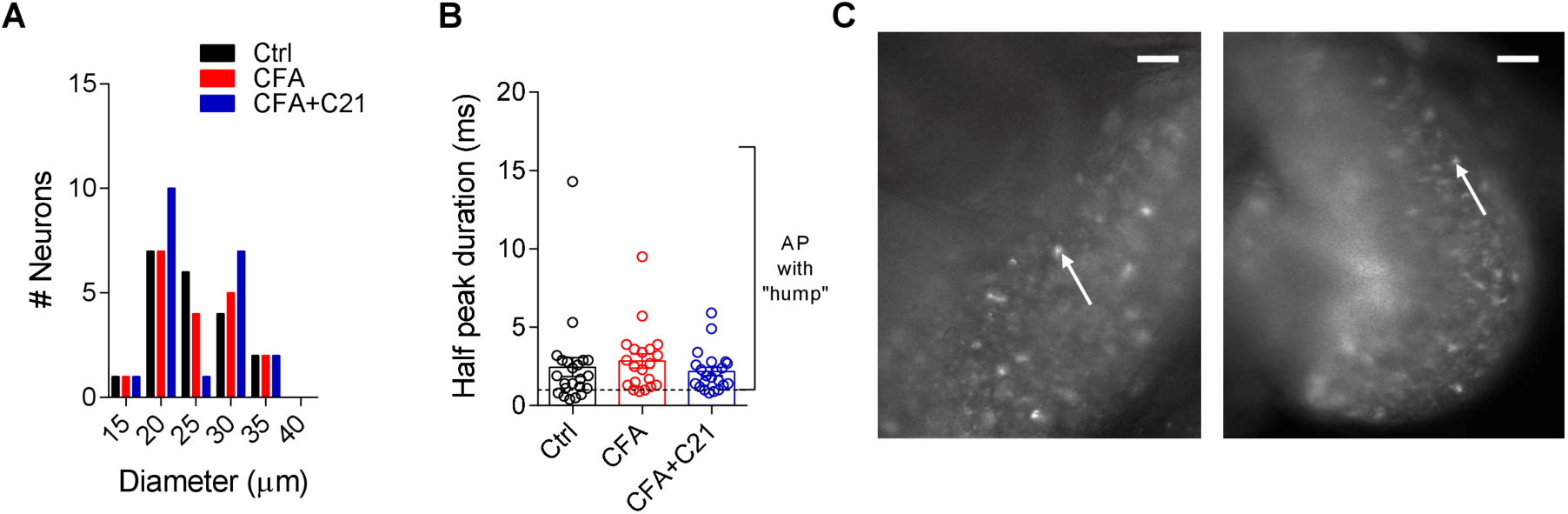
Post-hoc validation and neuronal profile of AAV-PHP.S-hSyn-G_i_DREADD transduced neurons. A) Distribution of neuronal diameters transduced by G_i_DREADD and characterized using whole cell patch clamp electrophysiology. Data from 2 female and 2 male mice B) Distribution of half peak duration (HPD) of action potentials (AP) recorded from transduced neurons. The black dotted horizontal line corresponds to HPD = 1 ms, thus demonstrating that most neurons had wide AP. C) Representative image of freshly dissected L3 DRG (left panel = CFA injected side, right panel = contralateral side) showing AAV-PHP.S-hSyn-G_i_DREADD labelled neuron (white arrows), scale bar = 250 μm

**Table S1:** Plasmids used for virus packaging.

| Plasmids | Addgene # | Acknowledgement |
| --- | --- | --- |
| pAAV-hSyn-hM4D(Gi)-mCherry | 50475 | Bryan Roth |
| pAAV-hSyn-hM3D(Gq)-mCherry | 50474 | Bryan Roth |
| pUCmini-iCAP-PHP.S | 103006 | Viviana Gradinaru |

## Notes

#### Summary of Updates

Amended text and updated supplementary figures with extra info.

